# A 4-pole Theory-Based Compensation Method using Multiple Loads for Impedance Spectroscopy of Porcine Brain Tissue

**DOI:** 10.1101/2025.11.19.689299

**Authors:** Moritz Gerber, Markus Symmank, Lucas Poßner, Michael Wallenta, Uwe Pliquett, Matthias Laukner, Thomas Knösche, Erdem Güresir, Florian Wilhelmy, Konstantin Weise

## Abstract

Accurate determination of the electrical properties of brain tissue—conductivity and relative permittivity—is essential for modeling and optimizing neurostimulation and neuroimaging and modulation techniques such as EEG, MEG, TMS, and TES. However, large discrepancies in reported conductivity values highlight the methodological challenges associated with impedance measurements. In this study, we present a systematic evaluation of compensation methods in impedance spectroscopy to improve measurement accuracy using a self-developed four-point platinum electrode designed for biological tissue applications. Three compensation algorithms—open-short, open-short-load, and multiple-load—were implemented and compared using an RC parallel test circuit that emulated typical measurement conditions, with load resistances ranging from 250 Ω to 5000 Ω. Quantitative error analysis demonstrated that the multiple-load method achieved the lowest normalized root mean square error (3.4%) and superior stability across frequencies from 100 Hz to 1 MHz. The calibrated probe was subsequently applied to post mortem porcine brain tissue, yielding conductivity values of grey matter of 0.109 S/m and white matter of 0.074 S/m. These results are significantly lower than standard simulation parameters of 0.275 S/m, and 0.126 S/m for grey and white matter, respectively. Finite-element simulations revealed that using the measured conductivities increased electric field predictions by 2.1% for TMS and 46% for TES in the maximum electric field magnitude, respectively. Our findings demonstrate that accurate impedance compensation is critical for reliable characterization of brain tissue and for enhancing the precision of neurostimulation modeling in the future.

## I. Introduction

Reliable and accurate measurements of the electrical material properties of brain tissue are essential because almost all imaging and stimulation techniques, such as electro-encephalography (EEG), magnetoencephalography (MEG), transcranial magnetic stimulation (TMS), or transcranial electrical stimulation (TES) require these material properties for modeling, simulation, and evaluation [1-3]. Studies have shown that there are notable variations in reported conductivity values for brain tissue [4-5]. This indicates that obtaining consistent and comparable measurements of electrical conductivity and relative permittivity of brain tissue presents methodological challenges. Numerous practical applications of electrical impedance spectroscopy (EIS) aim to evaluate relative differences between different tissues like healthy and pathologic tissue. However, the current challenge we want to address lies in the determination of accurate absolute values of the electrical material properties rather than relative differences. Therefore, we developed a 4-point platinum electrode, specifically designed to enable in vivo measurements using the four-point measurement method. The use of platinum was chosen due to its excellent biocompatibility, making it suitable for direct application in biological tissues in vivo. Besides numerous advantages, it is noted, that the 4-point method has still several disadvantages. First, parasitic cross impedances, like capacitances and conductances cannot be eliminated, second, inductive coupling from the current to the voltage channel cannot be avoided, and most important, the evaluated impedance has to be interpreted as a transversal (quasi) impedance because of the decoupling between the current and voltage channel. It is therefore essential to compensate the measured raw data during post-processing by using well known calibration data together with distinct compensation methods. Several compensation methods are available, e.g., *open-short, open-short-load*, and *multiple-load* method. In this work, we aim to systematically compare the aforementioned methods and to evaluate their applicability for EIS of brain tissue. This allows us to identify the strengths and limitations of each method in a controlled and comprehensible manner, providing a solid foundation for their evaluation in more complex experimental scenarios. Finally, the post-processed impedance data is used to determine equivalent circuit parameters of biological cell models, like the *β*-dispersion equivalent circuit model and the Cole-model [6].

## II. Methods

### A. Comparison of compensation methods

The test environment to compare the different compensation methods consists of an RC parallel circuit. The resistance represents the device under test (DUT) to measure and the capacitor ( *C* = 1.31 nF) represents the simplified parasitic properties of the measurement setup. The choice of resistance values is motivated by previous measurements of porcine brain, where ranges between 1000 Ω and 4000 Ω were observed using the 4-point probe described in [7]. To evaluate the performance of the compensation methods also outside of the measuring range, loads in the range from 250 Ω to 5000 Ω in steps of 250 Ω were investigated using the measurement setup described in [7]. After acquiring the recordings of the impedances, the compensation algorithms were applied to the raw data during post-processing. The equations for the *open-short* (OS), and the *open-short-load* (OSL) method, are derived from multipole theory [8]. Both methods yield the compensated impedance (*<u>Z</u>*_*c,OS*_ or *<u>Z</u>*_*c,OSL*_) and require reference impedances measured in a short circuit (*<u>Z</u>*_*s*_) and an open circuit (*<u>Z</u>*_*o*_). The *open-short-load* method utilizes an additional reference impedance with known impedance (*<u>Z</u>*_*l*_). The measured raw impedance is denoted by *<u>Z</u>*_*m*_ and the raw reference impedance is denoted by *<u>Z</u>*_*l,m*_.

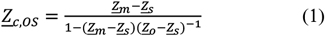

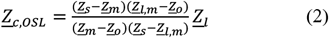

The equation for the *multiple-load* (ML) method can be derived from the multipole theory, by modulating the disturber as a 4-pole. In this case, we can describe the model with an inverted chain matrix (**[<u>A</u>]**^−1^), where *<u>U</u>*_*V*_ and *<u>I</u>*_*q*_ are the measured voltages and currents from the measurement device and *<u>U</u>*_*x*_ and *<u>I</u>*_*x*_ are the actual values of the DUT, and *a*_*ij*_ are the elements of the inverted chain matrix.

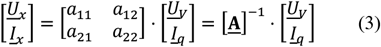

Dividing the voltage drop over the DUT ( *<u>U</u>*_*x*_), by the current flowing through the DUT (*<u>I</u>*_*x*_) yields the compensated impedance from the DUT (*<u>Z</u>*_*c,ML*_):

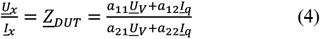

Dividing (4) by *a*_12_ and *<u>I</u>*_*q*_ yields the compensation equation for the *multiple-load* method:

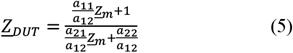

To be able to apply the compensation methods, one needs to determine the three ratios 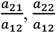 and 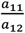 in (5). They can be determined by measuring the impedance of three reference loads. This leads to the following system of equations, where the load measurements are collected in the impedance matrix **[<u>Z</u>]** and the unknown ratios in the vector **[<u>a</u>]**. The matrix **[<u>Z</u>]** contains both the ideal resistances *<u>Z</u>*_*li*_ and the measured (raw) resistances *<u>Z</u>*_*li,m*_.

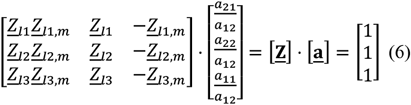

The unknowns in **[<u>a</u>]** can be determined by inverting the matrix **[<u>Z</u>]**:

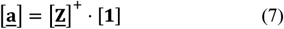

The error measures used to evaluate the performance of the compensation methods were the mean of the absolute error of the impedance 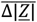, the mean of the absolute error for the phase angle 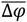, the median of the absolute percentage error (*MAPE*):

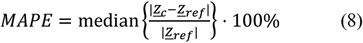

and the normalized root mean squared error (*NRMSE*) between the compensated and the reference data:

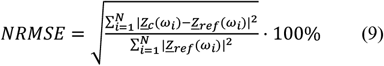

The errors were determined in the defined measuring range between 1000 and 4000 Ω and 100 Hz and 1 MHz.

### B. Probe

The 4-probe is manufactured by the Kurt-Schwabe Institute for Measurement and Sensor Technology Meinsberg e.V. (Waldheim, Germany).

The cable used for this application is a CAT 8.1 network cable, which has twisted and shielded pairs to reduce parasitic inductive effects.

The tip of the probe is guided out of the epoxy resin at an angle of approximately 25 degrees to enable its application during brain surgery. The electrodes assigned for current injection (I+ and I-) and voltage sensing (U+ and U-), have a diameter of 300 µm and are arranged in a square configuration with a spacing of approximately 550 µm, as shown in Fig. 1.

**Fig. 1.**
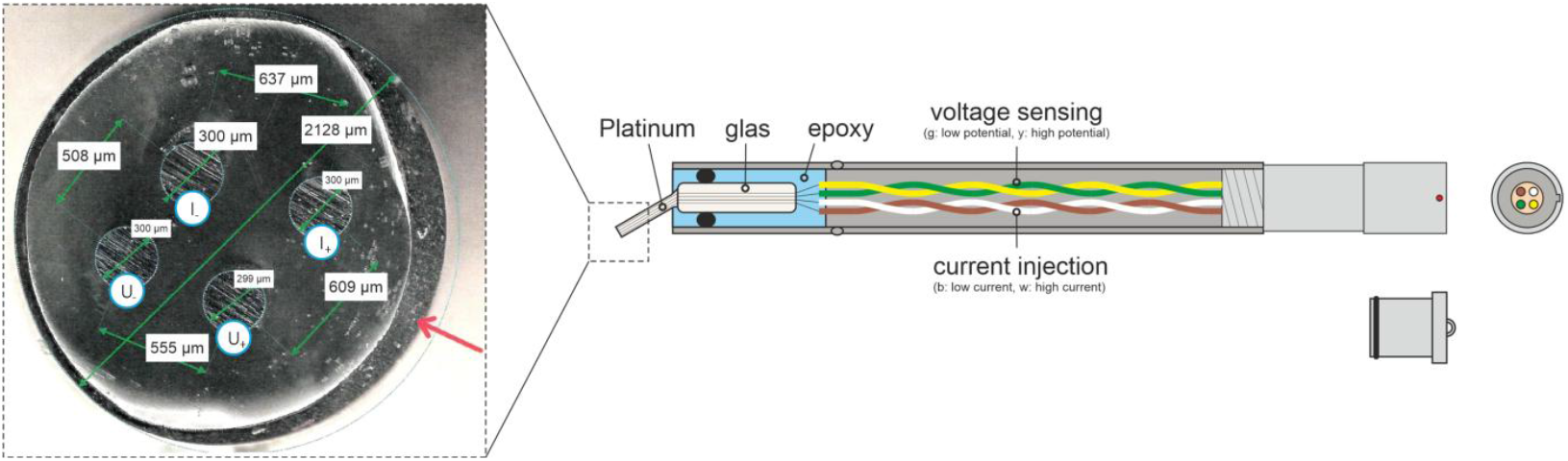
Microscopic and cross-sectional views of a multi-electrode probe designed for electrophysiological measurements. Left: Probe tip with four platinum electrodes embedded in a circular glass matrix, with diameters of 300 µm and a spacing of 508 µm to 637 µm. Right: Internal structure illustrating twisted wire pairs for current injection and voltage sensing, epoxy, and connector interface.

### C. Calibration Setup

The 4-point measurement reduces the influence of longitudinal impedance resulting from the properties of the probe or the double layers. Nevertheless, several limitations must be considered. Firstly, the technique measures transverse (quasi) impedance rather than true impedance. Secondly, capacitive parasitic effects can influence the measurement results. Therefore, it is essential to calibrate the probe using reference solutions with known conductivity. Solutions of potassium chloride (KCl) served as conductivity standards. Three calibrated solutions with conductivities of 147 μS/cm, 1,413 μS/cm, and 12,880 μS/cm were supplied by XS Instrument. Each solution was calibrated by the Danish Institute of Fundamental Metrology (DFM) and has a tolerance of 1%. A dilution series with a total of ten solutions was prepared. The conductivities of all solutions were measured using a conductivity probe (WTW TeraCon 325). All calibration measurements were performed in a thermal cabinet (VT 7011, Vötsch) set to 25 °C. The setup for calibration was identical to the setup used for the porcine brain measurement. The frequency response between 100 Hz and 1 MHz was measured at a current amplitude of 100 μA.

The impedance spectra were measured with the Medical Research ISX-3 analyzer from Sciospec Scientific Instruments GmbH (Bennewitz, Germany). The measurements were repeated *N* = 100 times for each solution. The uncompensated calibration data of the different solutions are shown in Fig. 2. In all solutions, the measured phase was close to zero at a frequency of 3 kHz. Therefore, at this frequency, the real part of the impedance can be used as a reference value for compensation. After compensating the raw impedance data <u>Z</u>of the solution data using the highlighted loads in Fig. 2 using (6) and (7), the geometry factor *k* is calculated by dividing the measured conductivity *σ* by the real part of the compensated admittance Re{*<u>Y</u>*_*c,ML*_} using the *multiple-load* method:

**Fig. 2.**
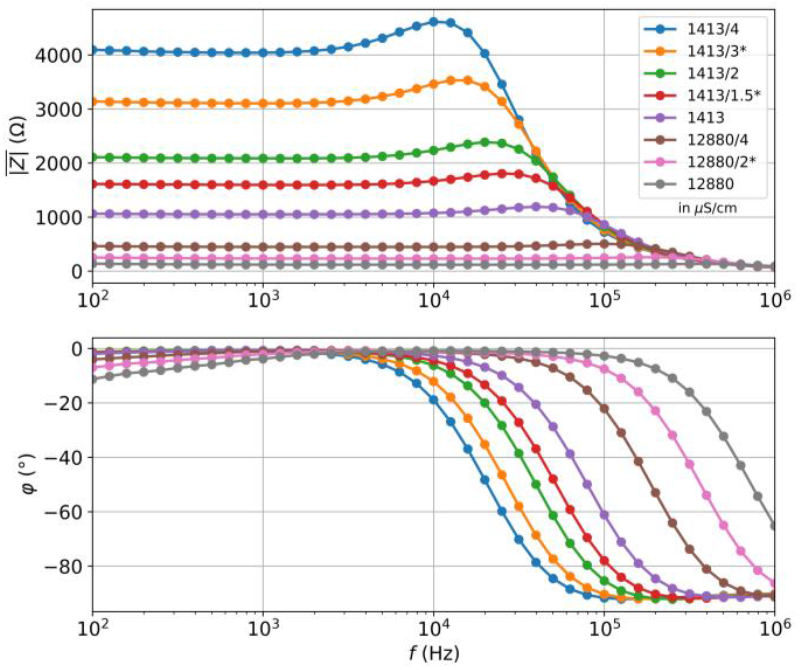
Raw calibration data acquired for potassium chloride (KCl) solutions of varying conductivity (μS/cm). The logarithmic plots show the mean impedance magnitude 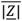 and mean phase angle 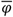 as functions of frequency f.

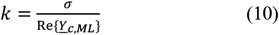

### D. Porcine brain measurement

To validate the measurement system, two porcine heads (*N* = 2) were obtained. Approximately 20 minutes elapsed between the time of death and the time of measurement for the first sample, and 70 minutes for the second sample. A craniotomy was performed on the heads to enable access for the measurements. The measurement protocol was tailored to align with the expected sterility standards of a surgical setting and were performed by a neurosurgeon. Multiple areas of the porcine brain were measured, e.g., the frontal lobe and the motor cortex.

The same measurement configuration was used as for the calibration, i.e. a current source with an amplitude of 100 µA at 41 logarithmically spaced frequencies between 100 Hz and 1 MHz. Each measurement took approximately one second. The probe was placed on different areas of the brain for three times (attempts). In each attempt, ten consecutive measurements (repetitions) were taken. Subsequently, the conductivity *σ* and the relative permittivity *ε*_*r*_ were calculated from the compensated measurement data using the geometry factor *k* determined during calibration:

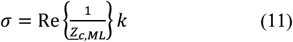

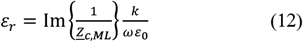

### E Fitting Data to Biological Tissue Models

For the quantitative characterization of biological tissues, the magnitude of the compensated impedance was fitted to two biological models. The aim is to derive tissue-specific parameters such as membrane capacitance and cell density from the impedance data. Model fitting allows a comparative evaluation of different tissues regarding their structural properties. To approximate the magnitude of the measured impedance (Fig. 3), the R–RC model and Cole-model were fitted using the Nelder-Mead optimization method. The R-RC model consists of the extracellular resistance ( *R*_*e*_), the membrane capacitance (*C*_*m*_), and the intracellular resistance (*R*_*i*_). The Cole-model extends this structure by including dispersion using a constant phase element (CPE), to capture non-ideal capacitive behavior. Specifically, the Cole-model comprises three components: a high-frequency resistance *R*_∞_, a residual low-frequency resistance *R*_0_, and a CPE with an impedance of 1/(*Q*_0_(*j*ω)^α^), where *Q*_0_ is a pseudo-capacitance, and α is a dispersion exponent characterizing the deviation from ideal capacitive behavior. An estimate of the cell density *P*_*y*_ of the measured cell package can be calculated from the Cole-model using the resistances *R*_∞_ and *R*_0_:

**Fig. 3.**
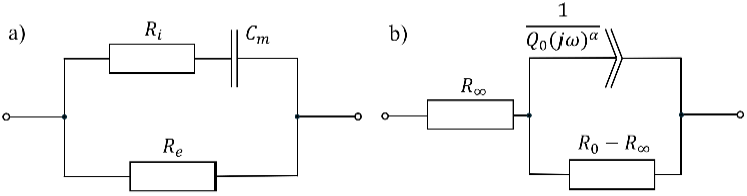
The R-RC and Cole models are equivalent circuit models used to fit impedance data from biological tissues. The R-RC model features parallel extracellular resistance and an RC branch for intracellular resistance and membrane capacitance. The Cole model adds the CPE.

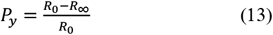

## III. Results

### A. Comparison of compensation methods

The uncompensated data for the artificial RC test circuit are shown in Fig. 4(a). The typical RC characteristics are clearly recognizable. Assuming that the resistors have a constant frequency response up to 1 MHz, the absolute value of the |*<u>Z</u>*| should remain constant across the entire frequency range at the respective resistance values, and the phase should be 0°. The results after applying compensation methods (OS, OSL, and ML) are shown in Fig. 2(b)-(d), and the corresponding error measures are given in Table 1. The *open-short* compensation performed worse, resulting in an NRMSE of 7.37%. Adding a load of 2500 Ω in the *open-short-load* compensation reduced the NRMSE by about half, to 3.4%. The *multiple-load* method outperforms the other compensation methods and the best results were obtained using three reference loads, which were chosen at symmetrical intervals. The error behavior observed in NRMSE is corroborated by similar patterns in MAPE and 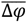. Increasing the number of reference loads led to higher residuals. This may be due to (i) limitations of the 2-port model in capturing parasitic effects, and (ii) the introduction of additional error sources, as the signal-to-noise ratio of the reference data is inherently limited. Therefore, the *multiple-load* method using three reference loads was used to compensate the impedance data of brain tissue.

**TABLE I.**
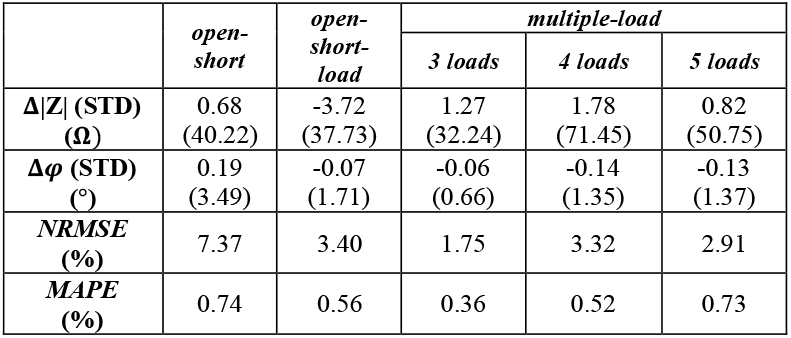
Evaluation of Compensation Methods with the rc-TEST CIRCUIT.

|  | *open-short* | *open-short-load* | *multiple-load* |  |  |
| --- | --- | --- | --- | --- | --- |
|  |  |  | 3 loads | 4 loads | 5 loads |
| $\Delta\vert Z\vert $ (STD) ($\Omega$) | 0.68<br>(40.22) | -3.72<br>(37.73) | 1.27<br>(32.24) | 1.78<br>(71.45) | 0.82<br>(50.75) |
| $\Delta\phi$ (STD) ($^\circ$) | 0.19<br>(3.49) | -0.07<br>(1.71) | -0.06<br>(0.66) | -0.14<br>(1.35) | -0.13<br>(1.37) |
| *NRMSE* (%) | 7.37 | 3.40 | 1.75 | 3.32 | 2.91 |
| *MAPE* (%) | 0.74 | 0.56 | 0.36 | 0.52 | 0.73 |

**Fig. 4.**
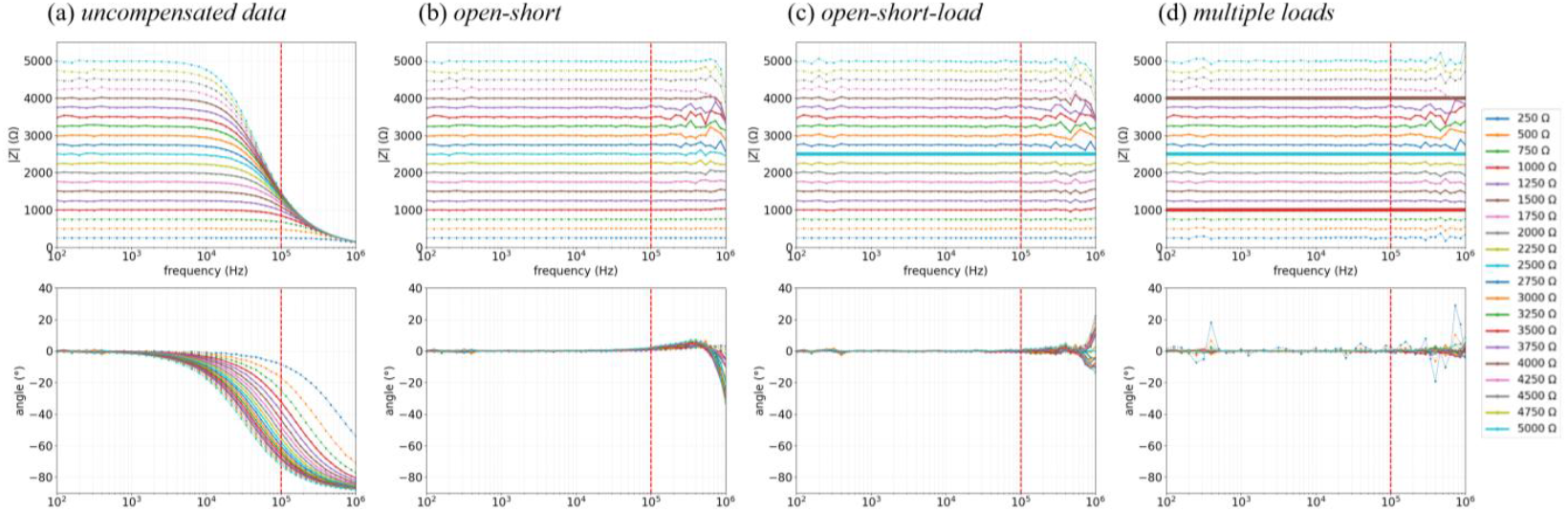
(a) The measurement of the uncompensated RC parallel test circuit; (b) *open-short* compensation; (c) *open-short-load* compensation with reference load at 2500 Ω; (d) *multiple-load* compensation using 3 loads at 1 kΩ, 2.5 kΩ, and 4 kΩ; (a-d) Thin dotted lines indicate measurements outside of the measuring range and thick lines in (c) and (d) indicate the reference loads used for compensation.

### B. Evaluation of calibration data

Different combinations of the calibration data shown in Fig. 2 were used as reference loads to identify the best possible combination for compensation. The error measure NRMSE between the compensation results and the ideal resistances were used to quantitatively assess the performance of each combination of references across the measured frequency range. The combination of 1413/3 µS/cm, 1413/1.5 µS/cm, and 12880/2 µS/cm, as shown in Fig. 5, yielded the lowest error and was therefore used for further analysis. The analysis of this combination also shows that effective compensation is possible with the integration of the probe. After compensation was performed, the parameter *k* = 150.86 1/m was determined using linear regression. The regression resulted in an *R*^2^ value of 0.999 (*p* ≪ 0.001) indicating superior linearity.

**Fig. 5.**
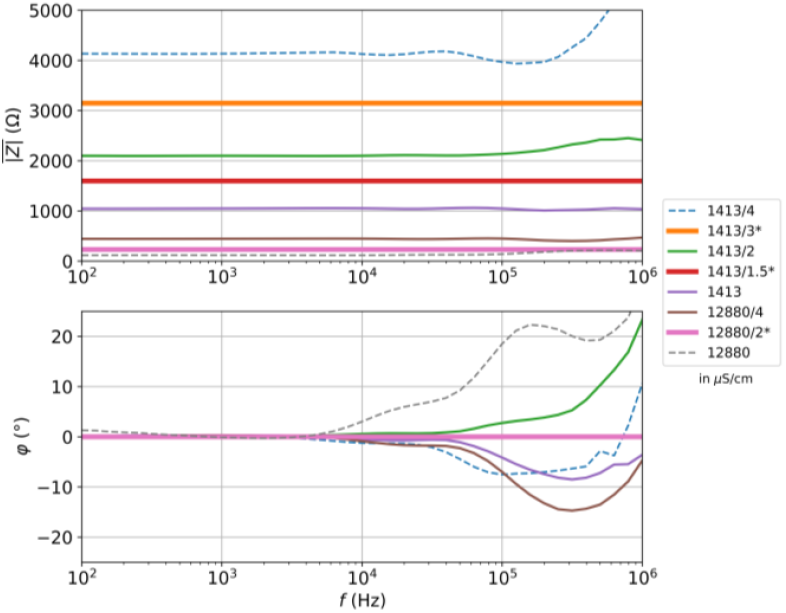
Frequency-dependent impedance magnitude |*<u>Z</u>*| and phase angle *φ* are shown for all conductivity conditions. Thick lines indicate the chosen references; dashed lines show compensated reference measurements outside the measurement range; solid lines represent compensated solutions within the measurement range.

### C. Porcine brain measurement

The porcine measurement data was categorized by brain region. As an example, Fig. 6 shows the compensated impedance magnitude |*<u>Z</u>*_*c,ML*_| measured from the motor cortex using the multiple-load method. The measurement data of the phase angle showed a considerable dispersion, so a joint optimization of magnitude and phase was not performed; instead, only the magnitude was used in the goal function for fitting. The fit of the R-RC model yielded a net membrane capacitance of *C*_*m*_ = 1.1 ⋅ 10^−9^ F for grey matter (GM) and *C*_*m*_ = 1.2 ⋅ 10^−9^ F for white matter (WM). By fitting the and *σ*_*WM*_ = 0.074 S/m for WM at a frequency of 3 kHz. Fig. 7 also shows the conductivity values determined for other areas of the brain. These were determined from measurements taken from two porcine brains using 3 attempts with 10 repetitions each. It is noted, that local differences can be identified.

**Fig. 6.**
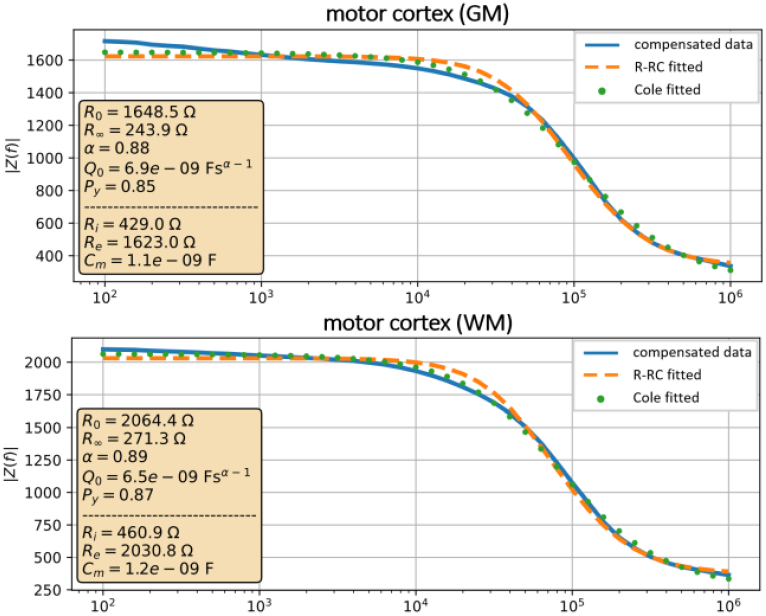
Impedance magnitudes |*<u>Z</u>*(*f*)| for grey and white matter in the motor cortex over a frequency range of 100 Hz to 1 MHz. The blue curve shows the compensated measurement data, while the orange and green curves represent the results of two model fits (R-RC model and Cole-model). The model parameters are listed in the respective legends.

**Fig. 7.**
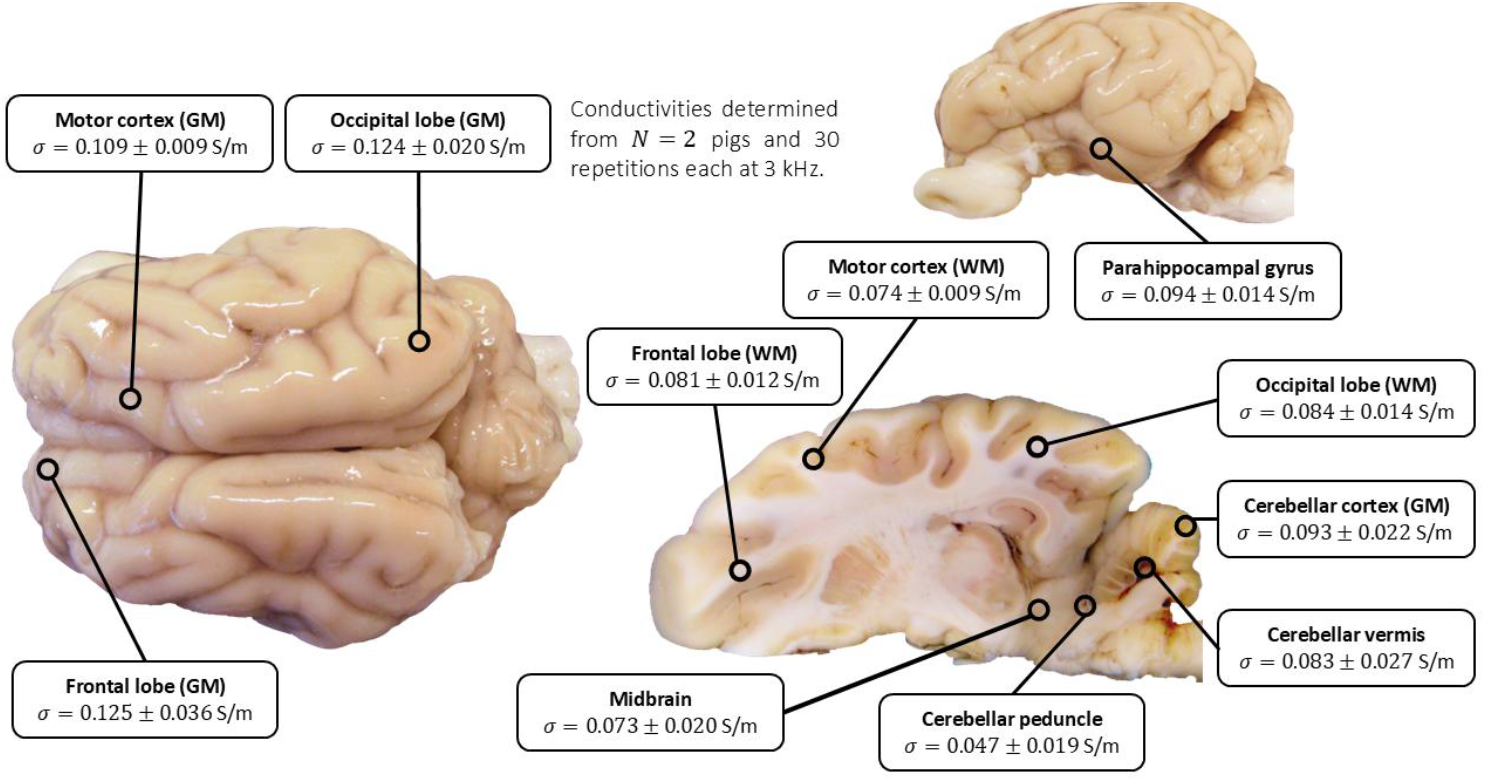
Regional electrical conductivities in S/m of different brain areas of a porcine, determined at a frequency of 3 kHz. Measurements were taken on grey and white matter as well as subcortical structures from 2 post mortem in situ from N = 2 animals with 30 repetitions per region. The anatomical representations of the brain are based on external image material taken from source [9].

## IV. Discussion

In the current study, we compared different compensation algorithms, namely the *open-short*, the *open-short-load*, and the *multiple-load* method in a simplified disturber setup using mostly idealized concentrated components. Theoretically, all methods can compensate for the disturbances ideally under the given 4-pole assumptions. However, we identified that the *multiple-load* method is superior compared to the other methods. This can be attributed to the technical difficulties involved in performing the necessary short and open measurements, as it requires the measurement of extremely low currents and voltages. It is noted that the transition from impedance to biological parameters by model fitting is only possible after compensation of the raw impedance data, which solidifies the importance of choosing a proper compensation method.

The major advantage of the *multiple-load* method is that the reference loads can be specifically adapted to the measuring range of the measuring devices and the measuring task. As a result, the currents and voltages to be measured are within a range that is well within the measuring capabilities of the devices and have a high signal-to-noise ratio. This motivated the application of the *multiple-load* method to the complex 4-point probe for compensating the measured data from brain tissue. The use of short and open measurements would in any case pose a difficulty in terms of practical implementation, as the contacts at the tip of the probe are very small, making electrical contact complicated.

It is important to note that the 4-pole approach used to derive the compensation equations is in fact only applicable to 2-point measurements. The electrical phenomena, i.e., the separation of the current and voltage measurements in a 4-point measurement setup, are not represented in the model. The phenomena that cannot be represented mainly occur in the upper frequency range above 100 kHz. For this reason, even in the idealized case, residual errors could still be observed after compensation when using concentrated components in the high frequency range. The remaining deviations in the upper frequency range are directly related to the fact that no “true” impedance can be determined from the 4-point measurement, but only a transfer (quasi) impedance, since the current and voltage measurements are decoupled from each other and their ratio has no physical meaning. To our knowledge, a mathematically exact compensation method based on multi-pole theory for 4-point measurements does not yet exist and is the subject of future research.

Previous studies demonstrate that uncertainties in tissue conductivity can substantially affect the accuracy of electric field modeling in non-invasive brain stimulation [10]. In the present study, we experimentally determined conductivity values for grey and white matter (*σ*_*GM*_ = 0.109 S/m, *σ*_*WM*_ = 0.074 S/m) that were considerably lower than the standard SimNIBS parameters (*σ*_*GM*_ = 0.275 S/m, *σ*_*WM*_ = 0.126 S/m).

To assess the impact of these differences, we conducted simulation studies for both transcranial magnetic stimulation (TMS) and TES targeting the motor cortex. For the TMS setup, the stimulation coil was positioned over the EEG electrode C3 and oriented along the central sulcus. In the TES condition, two rectangular electrodes (50×70 mm) were placed over C3 and AF4, respectively. Both configurations represent standard montages for the respective stimulation modalities. Subsequently, we compared the resulting electric fields for both conductivity models by evaluating the maximum electric field amplitude within the brain. For both TMS and TES, the use of our experimentally derived lower conductivity values led to an increase in the simulated electric field strength. The field enhancement was moderate for TMS (2.1%) but pronounced for TES (46%). This observation aligns with the findings of Saturnino et al. (2019), who also reported a greater sensitivity of TES to tissue conductivity variations compared to TMS. The discrepancy can be attributed to the different physical mechanisms underlying the two methods: whereas TMS induces electric fields primarily through time-varying magnetic flux, TES directly drives current through the tissues, making it inherently more dependent on local conductivity distributions.

Overall, our findings highlight the importance of using realistic, subject-specific conductivity estimates to improve the accuracy of electric field predictions.

## Acknowledgment

The project was greatly supported by Sciospec Scientific Instruments GmbH (04828 Bennewitz, Germany) providing rental equipment and Sensor Technology Meinsberg e.V. (04736 Waldheim, Germany) for the sensor design. We would also like to thank M. Dietrich (HTWK Leipzig) for his support in the assembly of the probe peripherals.

## Notes

### Competing Interest Statement

The authors have declared no competing interest.

## References

[1] G. B. Saturnino, A. Thielscher, K. H. Madsen, T. R. Knösche, and K. Weise, “A principled approach to conductivity uncertainty analysis in electric field calculations,” Neuroimage, vol. 188, pp. 821–834, 2019.

[2] A. V. Peterchev, T. A. Wagner, P. C. Miranda, M. A. Nitsche, W. Paulus, S. H. Lisanby, A. Pascual-Leone, and M. Bikson, “Fundamentals of transcranial electric and magnetic stimulation dose: definition, selection, and reporting practices,” Brain stimulation, vol. 5, no. 4, pp. 435–453, 2012.

[3] J. H. Cho, J. Vorwerk, C. H. Wolters, and T. R. Knösche, “Influence of the head model on EEG and MEG source connectivity analyses,” Neuroimage, vol. 110, pp. 60–77, 2015.

[4] T.R. Knösche and J. Haueisen, EEG/MEG Source Reconstruction. Cham: Springer, 2022.

[5] S. Gabriel, R. W. Lau, and C. Gabriel, “The dielectric properties of biological tissues: II. Measurements in the frequency range 10 Hz to 20 GHz,” Phys. Med. Biol., vol. 41, no. 11, pp. 2251–2269, Nov. 1996.

[6] U. Pliquett, “Bioimpedance: a review for food processing,” Food Engineering Reviews, vol. 2, no. 2, pp. 74–94, 2010.

[7] L. Poßner, F. Wilhelmy, J. Ranke, M. Wallenta, U. Pliquett, M. Laukner, E. Güresir, T. Knösche, and K. Weise, “Comparison of active and passive shielded 4-point probes for impedance spectroscopy of brain tissue,” in Proceedings of the 2024 International Workshop on Impedance Spectroscopy (IWIS), pp. 54–55, Sept. 2024, IEEE.

[8] Impedance Measurement Handbook; A Guide to Measurement Technology and Techniques; 6th Edition; 1990 (https://www.keysight.com/us/en/assets/7018-06840/application-notes/5950-3000.pdf, last access: June 7th 2025)

[9] Pig Brain – Amsterdam UMC Universitair Medische Centra; (https://anatomy-neurosciences.com/education/pigbrain/pig-outside-slices/, last access: November 11th 2025)

[10] G. B. Saturnino, A. Thielscher, K. H. Madsen, T. R. Knösche, and K. Weise, “A principled approach to conductivity uncertainty analysis in electric field calculations,” Neuroimage, vol. 188, pp. 821–834, 2019.

